# The receptor-like kinases BIR1 and BIR3 modulate antiviral resistance by different mechanisms

**DOI:** 10.1101/2025.01.24.634664

**Authors:** Carmen Robinson, Irene Guzmán-Benito, Ana Rocío Sede, Laura Elvira-González, Chenlei Hua, Malgorzata Ciska, Thorsten Nürnberger, Manfred Heinlein, César Llave

## Abstract

The BAK1-INTERACTING RECEPTOR-LIKE KINASE (BIR) proteins, including BIR1, are negative regulators of cell death and defense in Arabidopsis. Although BIR1 is upregulated during viral infections, its role in virus defense remains unclear. Here, we demonstrate a conserved role for the BIR family as modulators of antiviral resistance in Arabidopsis. During tobacco rattle virus (TRV) infection, *BIR* gene expression is regulated by antagonistic interactions between salicylic acid (SA) and jasmonic acid (JA) signaling pathways. Genetic evidence shows that BIR1, BIR2, and BIR3 negatively regulate TRV resistance via distinct mechanisms. The antiviral phenotype in *bir1-1* mutants is unaffected by mutations in the receptor-like kinases (RLK) BAK1, SOBIR1, or in the lipase-like PAD4. While BIR1 induction partially suppresses PTI marker gene expression, it does not affect early PTI responses such as reactive oxygen species (ROS) production or ethylene accumulation. BIR1 also inhibits plasmodesmata (PD) callose deposition and facilitates virus spread. Our data suggest that BIR1 modulates antiviral defense through mechanisms that include PTI gene expression, PD callose deposition alongside yet unidentified pathways independent of ROS or BAK1, SOBIR1, or PAD4 signaling components. In contrast, BIR3 suppresses an antiviral resistance pathway against TRV that requires BAK1 and the lipases PAD4 and EDS1. Our results suggest that BIR3 controls an NLR-dependent ETI-like response that restricts virus proliferation without triggering a hypersensitive response. These findings highlight the distinct roles of BIR family members in regulating antiviral defense in Arabidopsis.

## Introduction

Innate immunity in plants comprises the so-called pathogen-associated molecular pattern (PAMP)-triggered immunity (PTI) and effector-triggered immunity (ETI) (Jones *et al*., 2024). PTI prevents non-adapted microbes from infecting the host, and restricts infection of adapted pathogens in susceptible hosts, while ETI provides resistance against adapted pathogens (Zhou & Zhang, 2020). PTI uses cell surface receptors to sense immunogenic PAMPs, while ETI deploys nucleotide-binding domain leucine-rich repeat (LRR) intracellular immune receptors (NLRs) encoded by resistance (*R*) genes to sense pathogen-effector proteins or effector-induced manipulations of host proteins. NLRs are classified as TOLL/INTERLEUKIN RECEPTOR (TIR)-NLRs (TNLs) and coiled-coil domain containing NLRs (CNLs) (Saijo *et al*., 2018; Duxbury *et al*., 2021; Yu *et al*., 2021). Nevertheless, PTI and ETI function synergistically and interdependently in plants and share common downstream responses including the production of apoplastic reactive oxygen species (ROS), cytosolic calcium (Ca^+2^) influx, activation of mitogen-activated protein kinases (MAPK) and Ca^+2^-dependent protein kinases (CPK), generation of the signal molecule salicylic acid (SA), differential expression of genes, callose deposition and stomatal closure (Jones & Dangl, 2006; Ngou *et al*., 2021; Pruitt *et al*., 2021; Tian *et al*., 2021; Yuan *et al*., 2021). Besides, NLR-mediated resistance is often accompanied by a form of programmed cell death at the infection site known as hypersensitive response (HR) (Coll *et al*., 2011).

While RNA silencing has been traditionally regarded as the primary defense against viruses in plants (Voinnet, 2001), viral infections also activate canonical antimicrobial immune responses, such as mitogen-activated protein kinase (MAPK) activation, increased production of SA and ethylene (ET), callose deposition or the up-regulation of PTI-related genes (Korner *et al*., 2013; Nicaise, 2014; Calil & Fontes, 2017). Recently, growing evidence points towards an active role of several PTI and ETI signal transducers in the antiviral response. The receptor-like kinase (RLK) *BRASSINOSTEROID INSENSITIVE1-ASSOCIATED RECEPTOR KINASE 1* (*BAK1*), a member of the *SOMATIC EMBRYOGENESIS RECEPTOR KINASE* (*SERK*) family, is a positive regulator of several ligand-binding PTI receptors (Kemmerling *et al*., 2007; DeFalco & Zipfel, 2021). A role for *BAK1*, and its closest homolog *BAK1-LIKE1* (*BKK1*), in antiviral resistance has been reported using *bak1* and *bak1 bkk1* Arabidopsis mutants, which show increased susceptibility to several RNA viruses, including turnip crinkle virus (TCV), oilseed rape mosaic virus (ORMV), tobacco mosaic virus (TMV), plum pox virus (PPV) and tobacco rattle virus (TRV) (Yang *et al*., 2010; Korner *et al*., 2013; Nicaise & Candresse, 2017; Guzman-Benito *et al*., 2019). Interestingly, other RLK proteins interact with several viral proteins during infection (Fontes *et al*., 2004; Mariano *et al*., 2004; Li *et al*., 2018; Rosas-Diaz *et al*., 2018), although it does not necessarily imply interferences with PTI defense signaling cascades (Zeng *et al*., 2018; Macho & Lozano-Duran, 2019). *MAPK4* homologs in soybean (*Glycine max*) negatively regulate PTI signaling and SA accumulation, and inhibit defense against soybean mosaic virus (SMV) (Liu *et al*., 2011). Conversely, *MAPKKKK-*lll, which positively regulates cell-death signaling, plays an antiviral function during infection with beet black scorch virus (BBSV) in *Nicotiana benthamiana* (Gao *et al*., 2022).

Given that viruses are obligate intracellular biotrophs, the question arises as to how virus-derived PAMPs formed inside the cell can be recognized by host surface receptors to initiate a canonical PTI response. Interestingly, double-stranded RNA (dsRNA) triggers an array of PTI responses dependent on the co-receptor SERK1 (Niehl *et al*., 2016). Among them, dsRNA reduces plasmodesmata (PD) permeability by inducing the deposition of callose, thus restricting viral movement (Huang *et al*., 2023). *SERK1* and *BOTRYTIS INDUCED KINASE1* (*BIK1*) were identified as important signaling components required for dsRNA-induced PD callose deposition (Huang *et al*., 2023). Accordingly, dsRNA-induced protection of Arabidopsis plants against infection with ORMV is significantly reduced in *serk1* and *bik1 pbl1* double mutants (Niehl *et al*., 2016; Huang *et al*., 2023). While these findings demonstrated that dsRNA serves as an authentic elicitor of antiviral PTI, the specific ligand receptors involved in dsRNA perception remain elusive (Niehl & Heinlein, 2019).

ETI similar to that employed in response to nonviral pathogens can also be activated in plants upon recognition of viral effectors (Gouveia *et al*., 2016). Antiviral ETI involves the activation of intracellular NLR receptors to initiate a HR response that confines the virus to the infection site (Caplan *et al*., 2008; Zvereva & Pooggin, 2012). Correspondingly, several dominant antiviral *R* genes and their virus-encoded avirulent proteins have been reported in different plant-virus pathosystems (Piau & Schmitt-Keichinger, 2023). NLR-mediated signaling during ETI requires the combined activities of lipase-like proteins encoded by *ENHANCED DISEASE SUSCEPTIBILITY* (*EDS1*), *PHYTOALEXIN DEFICIENT 4* (*PAD4*) and *SENESCENCE ASSOCIATED GENE 101* (*SAG101*) (Sun *et al*., 2021). In Arabidopsis, these components are crucial for TNL-mediated HR and resistance against TCV underscoring the overlap between viral and non-viral signaling pathways (Zhu *et al*., 2011).

The plasma-membrane localized BAK1-INTERACTING RECEPTOR-LIKE KINASE1 (BIR1) represses antiviral defense in Arabidopsis since *bir1-1* mutants exhibit resistance against TRV (Guzman-Benito *et al*., 2019). BIR1 is one of the four members of the BIR family of RLKs in Arabidopsis, of which BIR1, BIR2 and BIR3 are negative regulators of PTI signaling (Gao *et al*., 2009; Halter *et al*., 2014; Imkampe *et al*., 2017). In the absence of elicitors, BIR proteins associate constitutively with BAK1, thereby preventing its interaction with ligand-binding receptors and the formation of functional BAK1-receptor complexes (Gao *et al*., 2009; Halter *et al*., 2014; Imkampe *et al*., 2017). Here we investigated the role of BIR proteins during viral infections. Our results indicate that the induced expression of *BIR* genes in virus-infected Arabidopsis involves the antagonistic interaction between the salicylic acid (SA) and jasmonic acid (JA) pathways. BIR1, BIR2 and BIR3 function as negative regulators of resistance against TRV, albeit by different mechanisms. Induction of BIR1 interfered with PTI gene expression and PD callose deposition triggered by different viral and microbial elicitors, whereas no effects on ROS accumulation or ET production were observed. Conversely, induction of BIR3, but not BIR1, represses an antiviral defense cascade requiring the PTI and ETI components BAK1, EDS1 and PAD4. These findings highlight the interplay between PTI and ETI signaling in regulating antiviral immunity and underscore the role of BIR proteins as key modulators of plant-virus interactions.

## Material and methods

### Plant material and transient expression assay

*Arabidopsis thaliana* Columbia-0 (Col-0) plants were grown in controlled environmental chambers under long-day conditions (16h day/8h night) at 22°C. Mutants used in this study included *sobir1-12, bir1-1 sobir1-1*, *bir1-1 pad4-1*, *bir1-1 eds1-2*, *bir1-1 bak1-4*, *bir1-1 pad4-1 sobir1-1*, *bir1-1 pad4-1 sobir7-1* (provided by Yuelin Zhang, University of British Columbia, Canada); *eds1-2*, *pad4-1* and *eds5-3* (from Jane E. Parker, Max Planck Institute for Plant Breeding Research, Germany); *bak1-4*, *bir2-1*, *bir3-2*, *csa1-2*, *bir3-2 bak1-4*, *bir3-2 bak1-4 csa1-2*, and *bir3-2 bak1-4 eds1-*12 (from Birgit Kemmerling, Center of Plant Molecular Biology, University of Tübingen, Germany). The dexamethasone (DEX)-inducible system for BIR1-mCherry gene expression and the generation of Arabidopsis Col-0 transgenic lines (BIR1 L6 and BIR1 L9) were described previously (Guzman-Benito *et al*., 2019). The Arabidopsis *BIR1* genomic sequence was PCR amplified and cloned into the pGWB14 binary vector, containing a C-terminal 3x human influenza hemagglutinin (HA) tag, using GATEWAY technology. The resulting HA-tagged BIR1 construct was driven by the CaMV35S promoter. Agrobacterium-mediated infiltration of *N. benthamiana* leaves was performed as described (Johansen & Carrington, 2001).

### Virus inoculation

Three weeks after germination, Arabidopsis plants were mechanically inoculated with sap from upper, non-inoculated leaves of *N. benthamiana* plants systemically infected with a TRV infectious clone tagged with green fluorescent protein (GFP) (Donaire *et al*., 2008). The sap was prepared by grinding infected tissue in 0.5M phosphate buffer (pH 7.0) at a 1:1 (W/V) ratio. Twelve microliters of the inoculum were gently rubbed onto the adaxial surface of three leaves per plant using carborundum as an abrasive. Relative TRV levels in individual Arabidopsis plants were measured 5 days post-inoculation (dpi), and data from plants infected with the same inoculum are presented within the same graph.

### Hormone treatments and determination of hormone content

Four-week-old Arabidopsis plants grown in soil were treated with 1 mM SA, 1 mM methyl jasmonate (MeJA) or 100 µM abscisic acid (ABA) in a solution containing 0.1% TritonX-100, and harvested after 3 h for gene expression analyses. Control plants were mock-treated with a solution containing 0.1% TritonX-100 and 0.01% EtOH. To measure JA and ABA content, rosette leaves from the same position were collected to minimize variability. Hormones were extracted and derivatized as described (Vallarino & Osorio, 2016), analyzed using gas chromatography coupled to time-of-flight mass spectrometry (GC-TOF-MS) (Pegasus III, Leco), and quantified using an internal standard ([2H4]-SA; OlChemIm Ltd, Olomouc, Czech Republic). For BIR1-mCherry induction, plants were sprayed daily for 3-4 days with 30 µM dexamethasone (DEX; Sigma) in a solution containing 0.02% of Silwet L-77 (Plant Media). Control plants were mock-sprayed with 0.02% of Silwet L-77 and 0.1% of EtOH. This treatment optimally induces the BIR1-mCherry transgene (Guzman-Benito *et al*., 2019).

### Quantitative RT-PCR assay

Total RNA was extracted using TRIzol reagent (Invitrogen) and treated with DNase I (Invitrogen) following the manufacturer’s instructions. One-step quantitative RT-PCR (RT-qPCR) was carried out using Brilliant III Ultra-Fast SYBR Green RT-QPCR Master Mix (Agilent Technologies) in a Rotor-Gene 6000/Rotor-Gene Q real-time PCR machine (Corbett/Qiagen) using a thermal protocol as follow: 48°C for 30 min; 95°C for 10 min; and 40 cycles of 95°C for 30 s, 64°C for 30 s, and 72°C for 20 s (Fernandez-Calvino *et al*., 2016). Relative viral accumulation and gene expression were determined using the delta-delta cycle threshold method and Rotor-Gene 6000 Series Software (Corbett). The housekeeping gene *CBP20* (*At5g44200*) was chosen for normalization because of its similar level of expression in mock-inoculated and TRV-infected tissues. Primers are listed in Table S1.

### Protein analysis

Leaf tissue samples were ground in liquid nitrogen and homogenized in extraction buffer (65 mM Tris-HCl, pH 8; 3% sodium dodecyl sulfate [SDS]; 1% ß-mercaptoethanol; 10% glycerol). Samples were diluted in Laemmli buffer, heated at 95°C for 5 min, and loaded onto 10% SDS-PAGE protein gels. Proteins were transferred to ECL nitrocellulose membranes (Amersham-Pharmacia) and detected using commercial horseradish peroxide (HRP)-conjugated secondary antibodies and a chemiluminescent substrate (LiteAblot Plus). Antibodies dilutions followed the manufacturer’s recommendations. Immunoprecipitation of mCherry-tagged BIR1 protein from Arabidopsis leaves was performed suing RFP-Trap (Chromotek) according to the manufacturer’s protocol.

### RNA and library preparation for RNA-Seq and bioinformatics data analysis

Total RNA was extracted using the RNeasy plant mini kit (Qiagen) and treated with DNase following the manufacturer’s instructions. RNA from six plants was pooled per sample, resulting in 12 samples (three biological replicates per treatment). RNA purity and concentration were assessed with a Nanodrop spectrophotometer (ThermoFisher Scientific), and integrity was confirmed using an Agilent Bioanalyzer 2100 system (Agilent Technologies). BIR1 transcript levels were measured via RT-qPCR. Sequencing libraries were prepared by NOVOGENE (HK) COMPANY LIMITED (Wan Chai, Hong Kong) (www.novogene.com) using NEBNext® UltraTM RNA Library Prep Kit for Illumina®. First-strand cDNA synthesis was done using random hexamer primer and M-MuLV Reverse Transcriptase (RNase H-), followed by second-strand synthesis with DNA Polymerase I and RNase H. After adaptor ligation and AMPure XP purification, libraries were amplified by PCR and quality-checked using the Agilent Bioanalyzer 2100 system. Sequencing was conducted on an Illumina PE150 Hiseq platform, generating paired-end reads.

Raw reads were filtered to remove adapters, poly-N regions, and low-quality sequences, yielding clean reads. These were mapped to the Arabidopsis TAIR v10 reference genome using HISAT2. Gene expression levels were quantified with HTSeq and normalized to Fragments Per Kilobase of transcript sequence per Millions of base-pairs sequenced (FPKM). Differentially expressed genes (DEGs) were identified using the DESeq2 R package (1.18.0), with *p-value*s adjusted for False Discovery Rate (FDR ≤0.05). Functional enrichment analysis of DEG, including Gene Ontology (GO) and KEGG, was performed using tGOseq and KOBAS, respectively, with significant enrichment set at corrected *p-value* ≤0.05. The raw sequencing data is available in the Gene Expression Omnibus under accession number GSE234036.

### Reactive oxygen species (ROS) and ethylene (ET) measurement

ROS and ET detection was conducted as described (Albert *et al*., 2015; Wan *et al*., 2019). Four[week[old BIR1 L9 or WT Arabidopsis plants were treated with 30 μM DEX (H_2_O as control) for 4 days at 24 h intervals. Leaves were cut into pieces and floated on H_2_O overnight before measurement. For ROS assays, leaf pieces were placed in a 96[well plate containing 20 μM L[012 (Wako Pure Chemical Industries Ltd) and 2 μg ml^-1^ peroxidase. Luminescence was measured using a Mithras LB 940 luminometer (Berthold Technologies) before (background) and for 1 h after flg22 (GenScript Biotech) or mock treatment. For ET determination, leaf pieces were incubated in 20 mM MES buffer (pH 5.6) with the specified elicitor (GenScript Biotech). ET accumulation was measured via gas chromatographic (GC[14A) by analyzing 1 ml of air drawn from the closed tube after the incubation period. All treatments were performed with at least three replicates.

### Callose staining

Callose staining was carried out as described (Huang *et al*., 2022; Huang *et al*., 2023). Three[week[old Arabidopsis BIR1 L9 or WT plants were sprayed with 30 μM DEX (H_2_O as control) for 4 days at 24 h intervals. Leaf disks were excised and floated on H_2_O. For *N. benthamiana* assays, leaves were agroinfiltrated with Agrobacterium cultures carrying a HA-tagged BIR1 construct or an empty vector (pGWB14) at OD_600_= 0.5. Samples were collected approximately 30 h post-infiltration and incubated in H_2_O for three hours. For callose staining, leaf disks were infiltrated with 0.1% aniline blue solution (50 mM potassium phosphate buffer, pH 9.0) containing 0.5 μg μl^-1^ poly(I:C) (Sigma) or 1 μM flg22. The disks were vacuum-infiltrated for 1-2 min (< 0.8 Pa), and imaged 30 min after treatment using Zeiss LSM 780 and LSM 980 confocal microscopes. Callose fluorescence was excited with a 405 nm diode laser, and mCherry fluorescence was excited at 561 nm, with emissions filtered at 475-525 nm, respectively. Callose fluorescence intensity was quantified with ImageJ using the calloseQuant plug-in (Huang *et al*., 2022). Callose spots were measured in 3-4 images per leaf disk, with regions of interest (ROIs) verified visually before analysis. Outliers (<1%) in fluorescence intensity were excluded from the analysis.

## Results

### SA and JA pathways antagonistically regulate BIR expression during viral infection

To investigate the role of BIR family members in viral infections, we examined their expression in Arabidopsis plants infected with TRV. Consistent with previous findings, RT-qPCR revealed that *BIR2* and, more significantly, *BIR3* transcripts were up-regulated in response to TRV infection compared to mock-treated plants (Figure 1a). *BIR4* transcripts were undetectable under our experimental conditions. Recent studies showed that TRV infection increases SA levels over time in Arabidopsis, and that *BIR1* transcription is SA-dependent during viral infections (Guzman-Benito *et al*., 2019). Here, we found that exogenous SA treatment also elevated *BIR2* and *BIR3* transcript levels, suggesting that their expression during the infection is also SA-dependent (Figure 1a).

**Figure 1.**
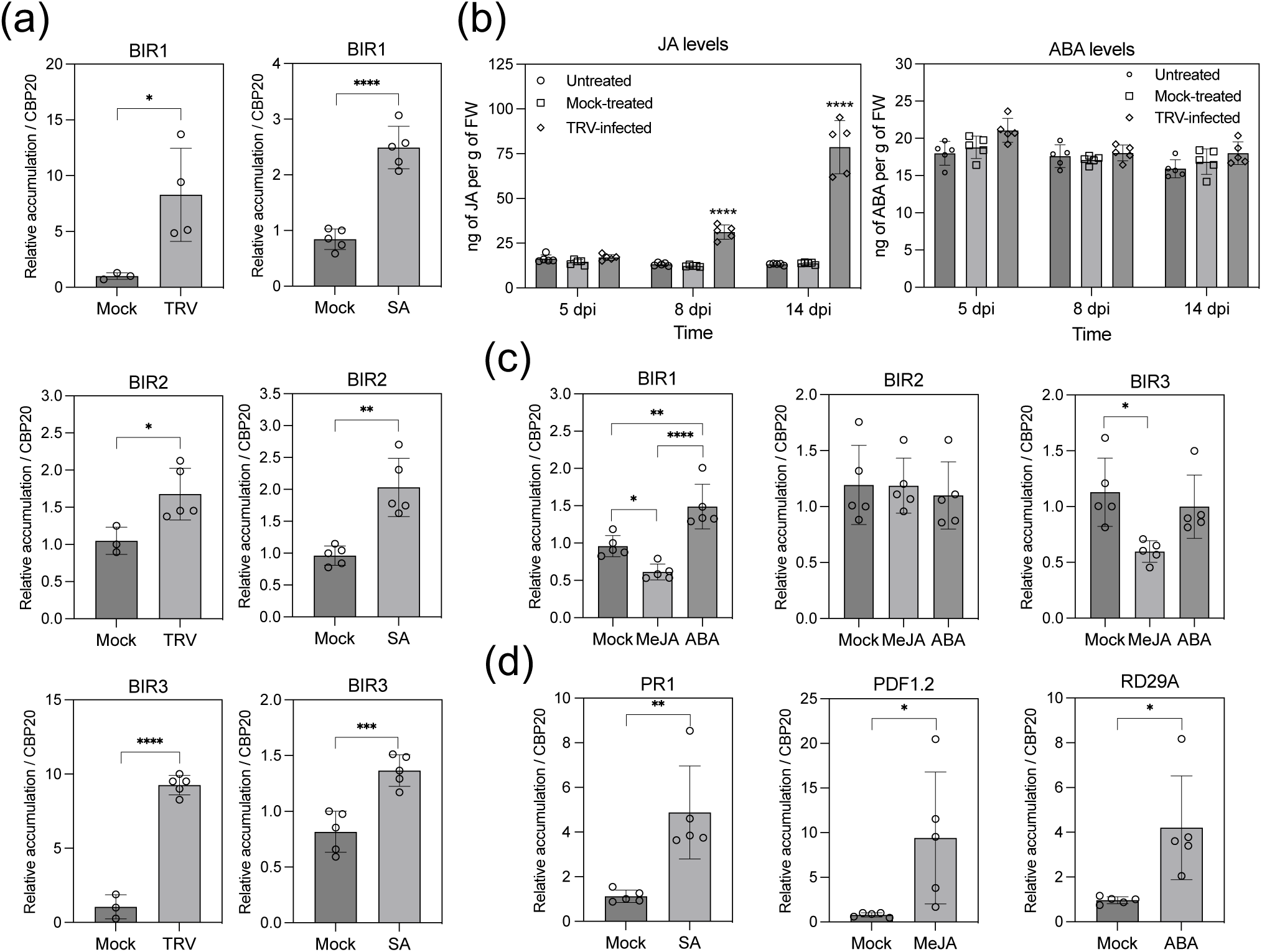
Hormone regulation of BIR gene expression in Arabidopsis. (a) RT-qPCR analysis of *BIR1*, *BIR2* and *BIR3* transcript levels in plants infected with tobacco rattle virus (TRV) (left) or treated with 1 mM salicylic acid (SA) (right) compared to mock-treated controls. (b) Time-course analysis of jasmonic acid (JA) and abscisic acid (ABA) levels in leaves of non-inoculated, mock-inoculated and TRV-infected Arabidopsis, determined by GC-TOF-MS. Error bars represent SD from five independent biological replicates. (c) RT-qPCR analysis of *BIR1*, *BIR2* and *BIR3* transcript levels wild type (Col-0) plants treated with 1 mM Methyl Jasmonate (MeJA) or 100 µM ABA. (d) RT-qPCR analysis of the hormone-responsive markers genes: *PR1* (SA), *PDF1.2* (JA) and *RD29A* (ABA) in response to SA, MeJA and ABA treatments. Measures were taken using the same set of replicates for each condition. All data were normalized to the *CBP20* internal control and compared to mock-treated plants (set to 1). Data are mean ± SD analyzed by unpaired *t-test* (a,d) or one-way ANOVA and Tukey’s multiple comparison test (b, c) ****, p <0.0001; ***, p< 0.001; **, p< 0.01; *, p <0.05. Experiments were repeated at least twice with similar results.

Interestingly, JA levels in Arabidopsis leaves gradually increased upon TRV infection compared to mock-treated or non-infected plants, indicating activation of both SA and JA pathways (Figure 1b). The interaction between SA and JA signaling is complex, with both synergistic and antagonistic effects depending on the pathogen (Zhao & Li, 2021). Notably, exogenous MeJA application suppressed *BIR1* and *BIR3* expression, while *BIR2* remained unaffected (Figure 1c). This suggests that the expression of *BIR1* and *BIR3* during infection is influenced by the antagonistic effects of JA and SA signaling.

Exogenous ABA treatment induced *BIR1* expression, but not *BIR2* or *BIR3* (Figure 1c). However, TRV-infected plants contained similar ABA levels to non-infected controls (Figure 1b), indicating a minimal role for ABA in *BIR* regulation during infection. RT-qPCR confirmed that SA, MeJA, and ABA treatments upregulated their respective marker genes, *PATHOGENESIS-RELATED 1* (*PR1*), *PLANT-DEFENSIN 1.2* (*PDF1.2*) and *RESPONSIVE*

*TO DESICCATION 29A* (*RD29A*) (Figure 1d). In conclusion, our data suggest an antagonistic regulation of *BIR* gene expression by SA and JA signaling during viral infection.

### BIR1 negatively modulates antiviral defense in Arabidopsis

The *bir1-1* mutants exhibit typical autoimmune phenotypes, including extensive cell death, constitutive activation of defense genes, increased SA levels, and resistance to the oomycete pathogen *Hyaloperonospora parasitica* Noco 2 (Gao *et al*., 2009). The *sobir1-1* mutation alleviates cell death and defense in *bir1-1* mutants, suggesting that the RLK *SUPPRESSOR OF BIR1-1 1* (*SOBIR1*) plays a role in *bir1-1* autoimmunity (Gao *et al*., 2009). In a previous study, we showed that *bir1-1 sobir1-1* plants exhibit antiviral resistance with reduced TRV levels compared to wild-type (WT) plants (Guzman-Benito *et al*., 2019). Here, we aimed to identify additional immune components involved in the antiviral response in *bir1-1* mutants. RT-qPCR confirmed that TRV levels at 5 dpi were significantly lower in *bir1-1 sobir1-1* mutants than in WT plants (Figure 2a). This antiviral effect was absent in *sobir1-13* single mutants, which accumulated TRV at levels similar to WT, indicating that the resistance in *bir1-1 sobir1-1* mutants is due to the loss of *BIR1* (Figure 2a). These results confirm that BIR1 negatively regulates an antiviral defense pathway independent of SOBIR1.

**Figure 2.**
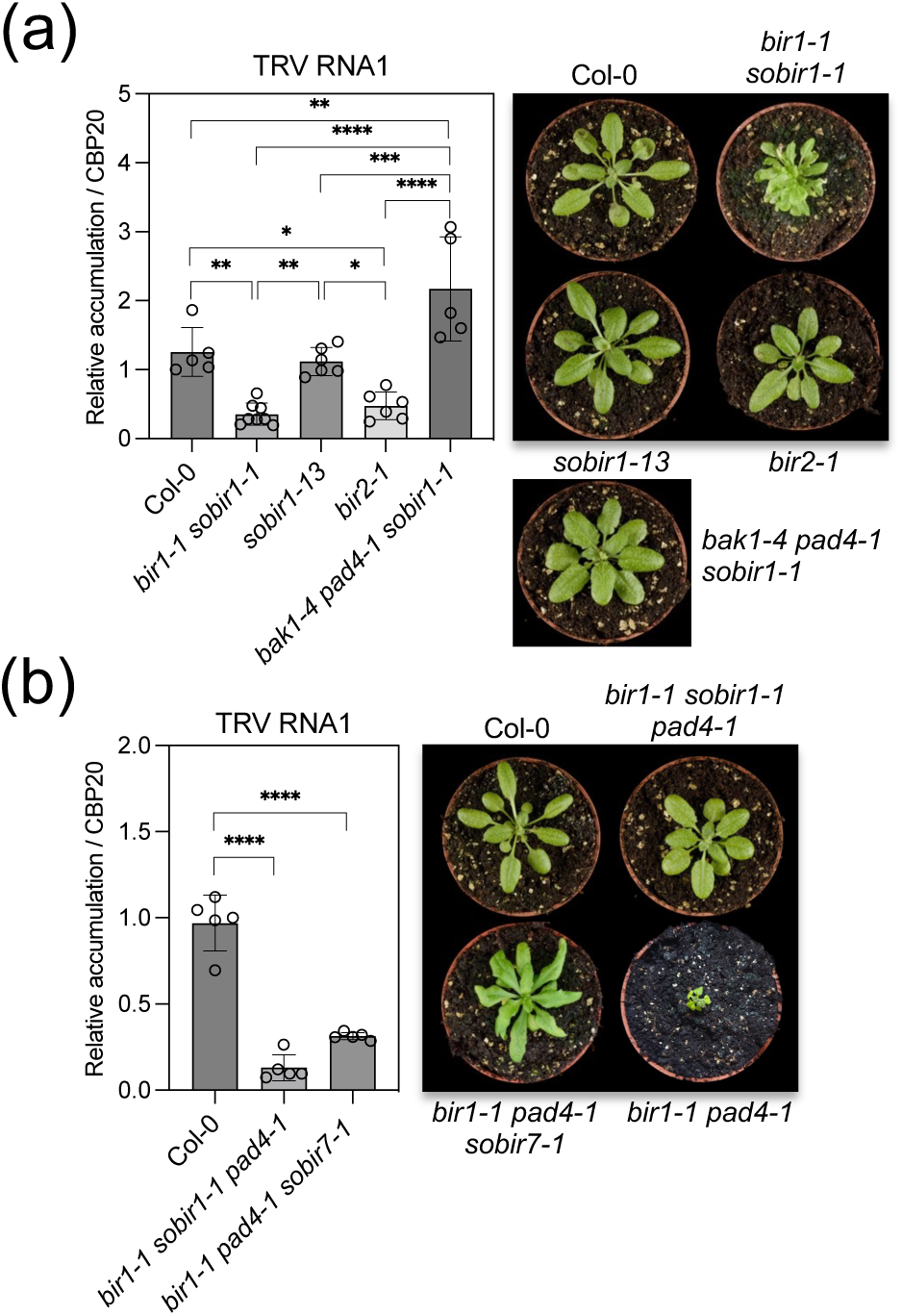
Genetic dissection of the antiviral response in *bir1-1* mutants. (a,b) RT-qPCR analysis of TRV genomic RNA levels in TRV-infected Arabidopsis wild-type (Col-0) and various mutant combinations, including *bir1-1 sobir1-1*, *sobir1-13*, *bir2-1*, *bak1-4 pad4-1 sobir7-1* (a), and *bir1-1 sobir1-1 pad4-1*, *bir1-1 pad4-1 sobir7-1* (b), at 5 days post-inoculation (dpi). Morphological phenotypes of each genotype at 5 dpi with TRV are also shown. A non-inoculated *bir1-1 pad4-1* mutant is included for size comparison. Relative TRV RNA levels were normalized to the *CBP20* internal control and compared to wild-type plants (set to 1). Data are presented as mean ± SD. Statistical significance was determined by one-way ANOVA followed by Tukey’s multiple comparison test; ****, p <0.0001; ***, p< 0.001; **, p< 0.01; *, p <0.05. Experiments were repeated at least twice with similar results

The autoimmune phenotype of *bir1-1* mutants also requires BAK1 and PAD4, as mutations in their genes suppress cell death and constitutive defense (Gao *et al*., 2009; Liu *et al*., 2016). To further explore the genetic basis of antiviral resistance in *bir1-1*, we examined the effects of *bak1-4* or *pad4-1* mutations on TRV accumulation in the *bir1-1* background. Arabidopsis double mutants *bir1-1 pad4-1*, *bir1-1 bak1-4* and *bir1-1 eds1-2*, due to their extreme dwarfism, suffered severe damage upon TRV inoculation, yielding inconsistent results. Therefore, we used the *bir1-1 sobir1-1 pad4-1* mutant, which displays WT morphology due to suppression of dwarfism by the *sobir1-1* mutation (Gao *et al*., 2009). This triple mutant accumulated significantly less TRV than WT at 5 dpi, suggesting that the antiviral phenotype of *bir1-1* is not abolished by the *pad4-1* mutation (Figure 2b). The constitutive defense and dwarfism in *bir1-1 pad4-1* were also restored by the *sobir7-1* allele, which carries a point mutation in *BAK1* (Liu *et al*., 2016). TRV accumulation was reduced in the *bir1-1 pad4-1 sobir7-1* triple mutant, indicating that the defense signaling in *BIR1*-defective plants is independent of *BAK1* (Figure 2b). In conclusion, our data show that BIR1 negatively regulates an immune antiviral response that limits TRV proliferation, independent of *SOBIR1*, *BAK1* and *PAD4*.

### BIR1 induction does not influence ROS production or ET accumulation during PTI

To investigate defense outputs downstream of BIR1 that could be eventually activated during viral infection, we examined how BIR1 transgenic induction affects key PTI signaling events. Reactive oxygen species (ROS) play a dual role in plant-pathogen interactions by promoting cell death in infected tissues and serving as signals for defense activation (Dat *et al*., 2000). A luminol-based assay showed sustained apoplastic ROS production in upper, non-inoculated leaves of *N. benthamiana* infected with TRV-GFP, compared to mock-treated controls (Figure 3a). This tissue accumulated high TRV-GFP levels (Figure 3b), suggesting that virus-derived GFP fluorescence might contribute to the background signal. As a control, a transient ROS peak was observed in response to flg22 in *N. benthamiana* leaf disks (Figure 3a).

**Figure 3.**
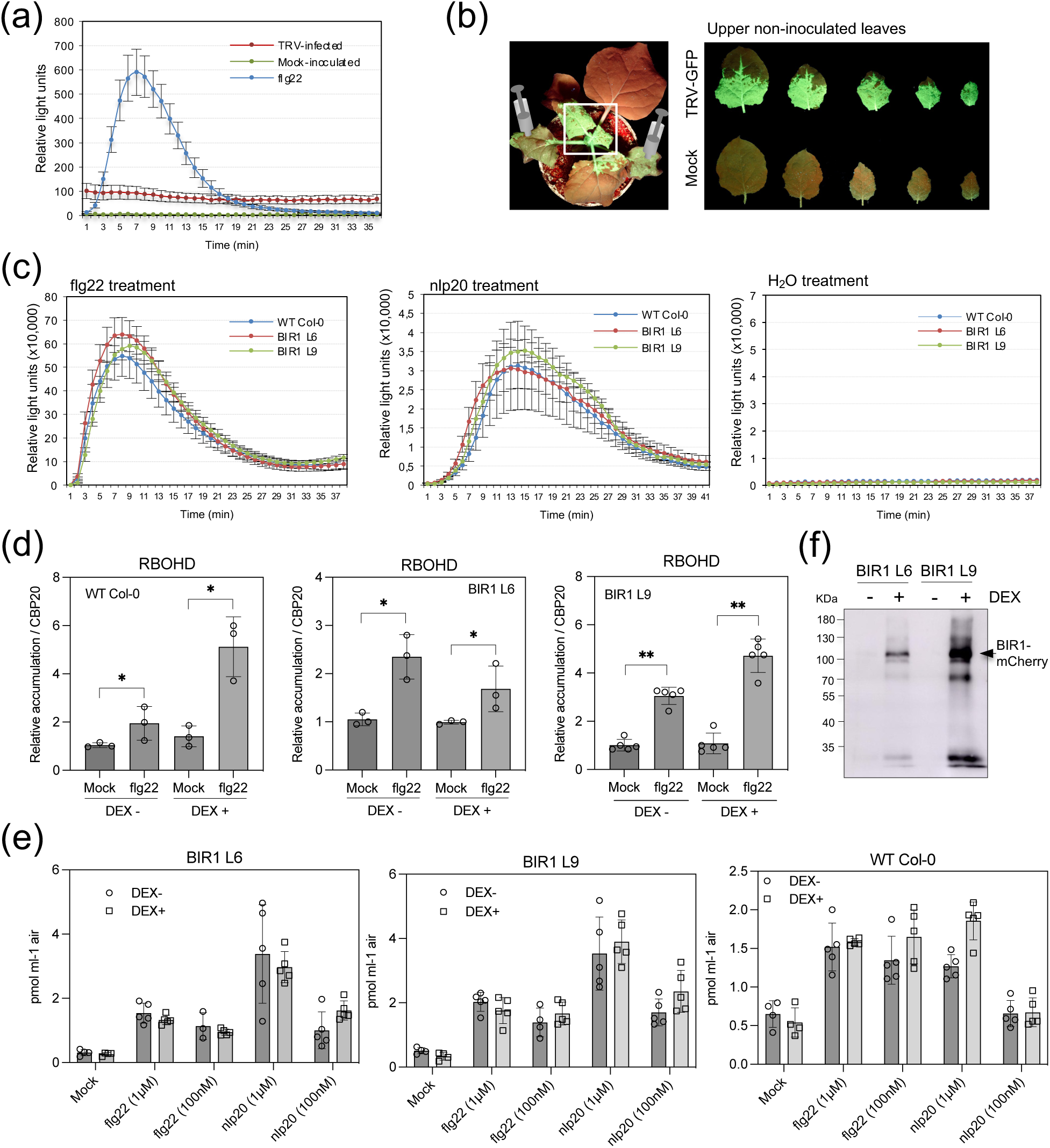
Effects of BIR1 induction on flg22- and nlp20-triggered PTI responses. (a) Time-course of reactive oxygen species (ROS) production in upper leaves of *N. benthamiana* systemically infected with an infectious clone of Tobacco rattle virus tagged with green fluorescent protein (TRV-GFP). Mock-inoculated plants and plants treated with 1 µM flg22 serve as controls. (b) Viral spread in young, non-inoculated leaves of *N. benthamiana* plants infected TRV-GFP. (c) ROS accumulation in dexamethasone (DEX)-treated wild-type (WT), BIR1 L6, and BIR1 L9 inducible plants after elicitation with 1 µM flg22, 1 µM nlp20, or H_2_O (mock). Data represent mean ± SD of relative fluorescence units. (d) RT-qPCR analysis of *RBOHD* transcript levels after treatment with H_2_O (mock) or 1 µM flg22 for 45 minutes in DEX-treated (DEX+) and non-treated (DEX-) samples. Values were normalized to the *CBP20* internal control and compared to mock-treated plants (set to 1). Data are presented as mean ± SD. Statistical significance was determined by unpaired t-test. **p < 0.01; *p < 0.05. (e) Ethylene accumulation in WT, BIR1 L6, and BIR1 L9 transgenic plants after treatment with H_2_O (mock), flg22, and nlp20 at 1 µM or 100 nM. Data represent average values ± SD. (f) Immunoprecipitation of mCherry-BIR1 fusion proteins (arrowhead) from DEX-treated and mock-treated samples of BIR1 L6 and BIR1 L9 plants used in (c) and (d). Experiments were repeated three times with similar results.

Next, we evaluated the impact of BIR1 induction on ROS accumulation using a dexamethasone (DEX)-inducible system to express mCherry-tagged BIR1 (Guzman-Benito *et al*., 2019). This approach enabled BIR1 activation without viral infection, avoiding confounding effects. BIR1 was induced in two transgenic lines (BIR1 L6 and BIR1 L9) with DEX, while controls received mock treatment. Leaf disks were incubated with flg22 or the nlp20 fragment of NECROSIS AND ETHYLENE-INDUCING PEPTIDE 1 (NLP) as elicitors. Flg22 and nlp20 were chosen because they activate immune responses through cell surface RLK-type pattern-recognition receptors (PRR) and SOBIR1-dependent RLP-type PRR, respectively (Ranf, 2017). We found that flg22 and nlp20 triggered a peak in apoplastic ROS, comparable across DEX-treated WT, BIR1 L6, and BIR1 L9 plants, indicating that BIR1 induction does not affect PTI-induced ROS accumulation (Figure 3c). We then measured RBOHD gene expression, which encodes the plasma membrane NADPH oxidase responsible for most ROS production during pathogen responses (Morales *et al*., 2016). RBOHD transcripts were elevated by flg22 treatment in both WT and BIR1 lines, regardless of DEX treatment, showing that BIR1 induction does not impair RBOHD up-regulation (Figure 3d). These results confirm that BIR1 induction does not affect the early oxidative burst triggered by PTI elicitors (Liu *et al*., 2016).

Previous research indicated that the cytosolic *ACETYL-CoA CARBOXYLASE I* (*ACC1*) gene is down-regulated in TRV-infected Arabidopsis, with the *acc1-1* mutant being more susceptible to TRV (Fernandez-Calvino *et al*., 2014). ACC1 is involved in the biosynthesis of very long-chain fatty acids (VLCFA), which activate ET biosynthesis. Although the antiviral role of ET is unclear (Zhao & Li, 2021), ET may be important in the defense against TRV. Thus, we hypothesize that BIR1 could influence ET production. In response to flg22 and nlp20 at concentrations of 1 µM and 100 nM, we observed similar ET levels in WT, BIR1 L6, and BIR1 L9 plants, regardless of DEX treatment, suggesting that BIR1 induction does not affect ET accumulation (Figure 3e). Accumulation of BIR1-mCherry protein in the BIR1 L6 and BIR1 L9 lines was confirmed by immunocapture followed by western blot (Figure 3f).

### BIR1 induction minimally affects the Arabidopsis transcriptome

To investigate if BIR1 induction affects the expression of defense genes in the absence of immune elicitors we studied the Arabidopsis transcriptome using the inducible transgenic line BIR1 L9. Three-week-old BIR1 L9 plants were treated with DEX for four days and harvested for RNA sequencing, with mock-treated plants as controls. Three biological replicates were analyzed for each condition: DEX-treated WT (DeWT_), mock-treated WT (MoWT_), DEX-treated BIR1 L9 (DeL9_), and mock-treated BIR1 L9 (MoL9_) plants (Table S2). After four days, BIR1 L9 plants showed WT morphology and accumulated BIR1 transcript and protein at levels similar to those observed during virus infection or SA treatments (Figure 4a,b) (Guzman-Benito *et al*., 2019).

**Figure 4.**
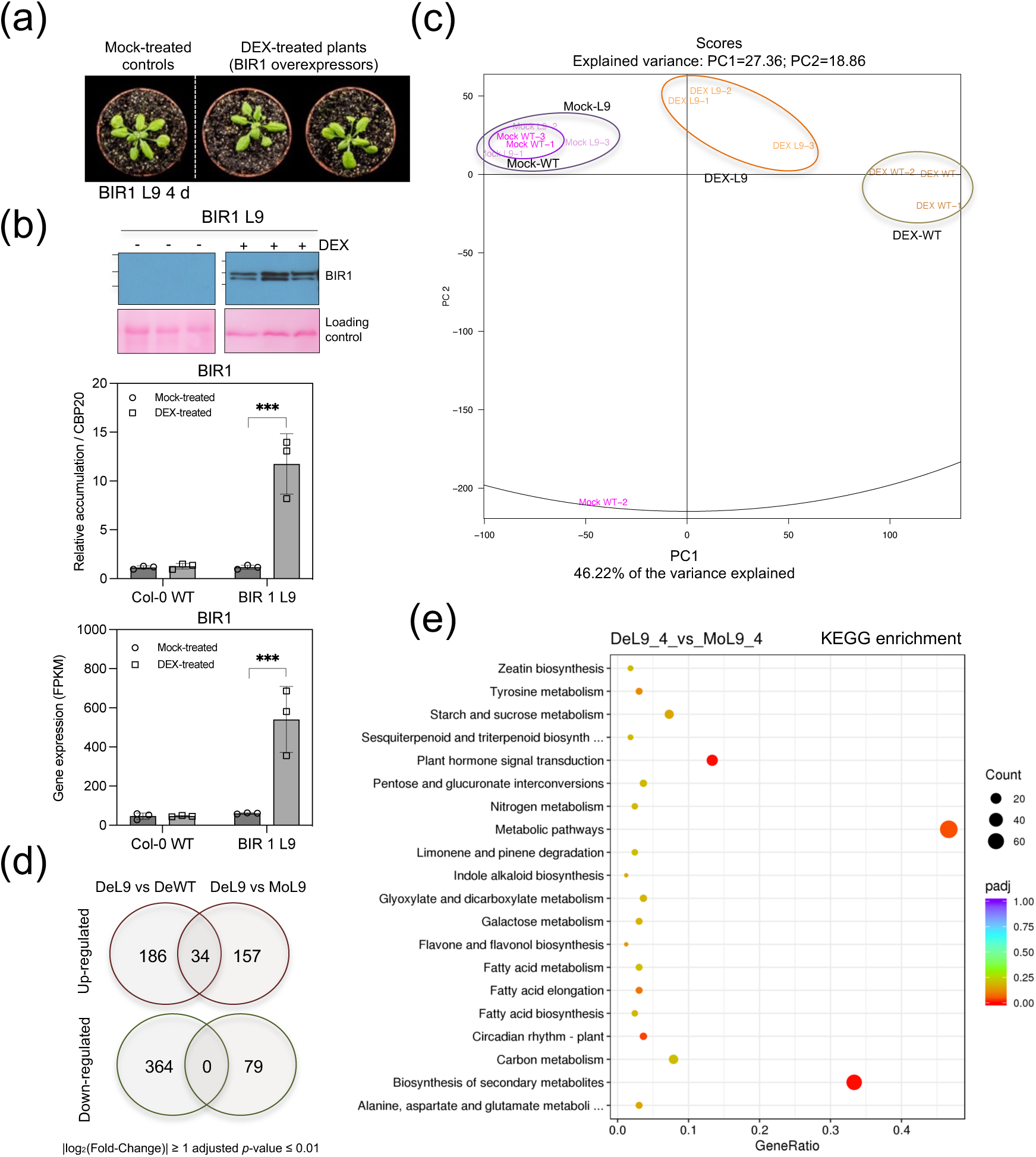
Transcriptional changes associated with BIR1 induction in Arabidopsis. (a) Morphology of representative dexamethasone (DEX)-inducible BIR1-mCherry transgenic Arabidopsis plants (line 9, L9) after four days of 30 µM DEX or mock treatment (H_2_O). (b) Western blot analysis using anti-mCherry antibody to detect BIR1-mCherry protein accumulation after four days of DEX treatment. BIR1 transcript levels in wild-type (WT) and BIR1 L9 plants were quantified using RNA-Seq (FPKM) and RT-qPCR. RT-qPCR values were normalized to the *CBP20* internal control, and compared to mock-treated (set to 1). (c) Principal component analysis (PCA) of RNA-Seq data from mock-treated WT (Mock-WT), mock-treated BIR1 L9 (Mock-L9), DEX-treated WT (DEX-WT), and DEX-treated BIR1 L9 (DEX-L9) plants. (d) Number of up- or down-regulated transcripts (|log_2_(Fold-Change)| ≥ 1 and adjusted p-value ≤ 0.01) in each pairwise comparison. (e) Scatterplot of selected KEGG terms of differentially expressed genes (DEGs) between DEX-treated and mock-treated BIR1 L9 plants. Dot size represents the gene count, and color indicates the significance of term enrichment. GO terms with adjusted p-value < 0.05 are significantly enriched. RT-qPCR data are presented as mean ± SD. Statistical significance was determined by unpaired t-test. ***p < 0.001. All experiments were repeated at least twice with similar results.

Principal component analysis (PCA) revealed consistency in the transcriptomic data across samples, with the three major clusters corresponding to DEX-treated BIR1 L9, DEX-treated WT, and mock-treated WT and BIR1 L9 samples (Figure 4c). The Pearson pairwise correlation coefficients confirmed high correlations within conditions (R² > 0.96) and across conditions (R² from 0.92 to 0.98) (Table S3). The PCA plot showed that DEX treatment caused differences between samples, but BIR1 induction had minimal impact on global gene expression (Figure 4c). Differentially expressed genes (DEGs) from various comparisons are shown in Figure 4d and Tables S4 and S5. Gene ontology (GO) and Kyoto Encyclopedia of Genes and Genomes (KEGG) pathway analysis revealed enrichment of genes involved in oxidoreductase activity, transmembrane transport, lipid and carbohydrate metabolism in both WT and BIR1 L9 plants (Figure 4e, Tables S6 and S7). These changes were predominantly observed in DEX-treated plants, indicating DEX-dependent effects (Figure S1). Importantly, RNA-Seq analysis showed that BIR1 induction or DEX treatment did not significantly misregulate genes involved in defense pathways during plant-pathogen interactions.

### Induction of BIR1 compromises the expression of defense genes during PTI

To identify defense mechanisms activated in *bir1-1* mutants potentially contributing to antiviral resistance against TRV, we examined whether BIR1 induction influenced the expression of defense genes in response to a PTI elicitor in Arabidopsis. To test this idea, plants from two independent inducible BIR1 transgenic lines (BIR1 L6 and BIR1 L9) were sprayed with DEX to induce BIR1 or mock-treated with H_2_O. Transcript levels of PTI marker genes *FLG22-INDUCED RECEPTOR-LIKE KINASE 1* (*FRK1*) and *PR1* were then measured following treatment with 1 µM flg22, used as a PTI elicitor. In the absence of DEX, both genes were significantly upregulated in flg22-treated but not H_2_O-treated WT, BIR1 L6, and BIR1 L9 plants (Figure 5a). In DEX-treated WT plants, *FRK1* and PR1 were similarly upregulated by flg22, ruling out any repressive effects of DEX on PTI gene expression under these conditions (Figure 5a). However, BIR1 induction in DEX-treated BIR1 L6 and L9 plants notably reduced flg22-induced expression of *FRK1* and *PR1* (Figure 5a). The induction of *BIR1* in these lines was confirmed by RT-qPCR (Figure 5b). In summary, BIR1 induction suppresses the flg22-elicited expression of PTI marker genes *FRK1* and *PR*1, suggesting that, upon PTI activation, BIR1 inhibits a key step in the signaling pathway leading to defense gene expression.

**Figure 5.**
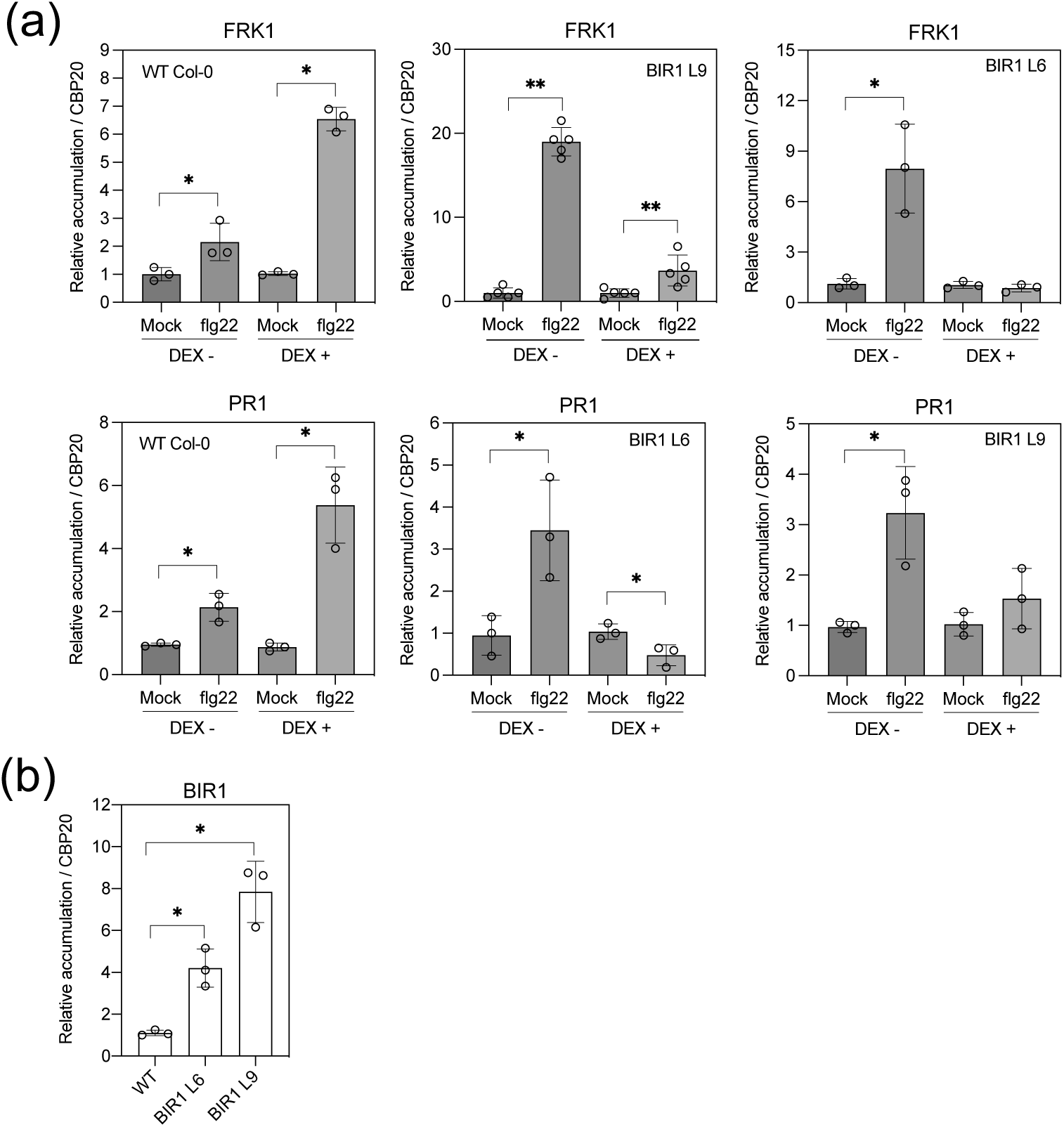
BIR1 induction interferes with flg22-triggered PTI gene expression. (a) RT-qPCR analysis of the PTI marker genes *FRK1* and *PR1* in wild-type (Col-0), BIR1 L6, and BIR1 L9 inducible plants after treatment with H_2_O (mock) or 1 µM flg22 for 45 minutes. Plants were pre-treated with or without dexamethasone (DEX) as indicated. (b) RT-qPCR analysis of *BIR1* transcript levels in DEX-treated samples prior to flg22 treatment. All data were normalized to the *CBP20* internal control and compared to mock-treated plants (set to 1). Data are presented as mean ± SD. Statistical significance was determined by unpaired t-test. **p < 0.01; *p < 0.05. All experiments were repeated at least twice with similar results.

### Transient expression of BIR1 inhibits PD callose deposition triggered by dsRNA and supports TRV spreading in *N. benthamiana*

Callose (β-1,3-glucan) deposition at PD regulates symplastic transport between plant cells (Wu *et al*., 2018). During viral infections, this process restricts viral movement between plant cells and acts as an early antiviral defense triggered by dsRNA replicative intermediates (Beffa *et al*., 1996; Huang *et al*., 2023). Viral movement proteins (MP) counteract this by suppressing dsRNA-induced callose deposition, promoting viral spread (Huang *et al*., 2023). To examine whether BIR1 expression inhibits PD callose deposition, we performed *in vivo* aniline blue staining to quantify PD-associated callose in *N. benthamiana*. Using the Agrobacterium-mediated transient assay, we expressed HA-tagged Arabidopsis BIR1 under the 35S promoter, with an empty vector (pGWB14/EV) as a control. We assessed callose deposition triggered by the synthetic dsRNA analog polyinosinic-polycytidylic acid [poly(I:C)] (0.5 µg µl^-1^) as described previously (Huang *et al*., 2023). We observed increased PD-associated callose levels in poly(I:C)-treated leaves infiltrated with EV compared to H_2_O-treated leaves (Figure 6a,b). However, poly(I:C)-induced callose deposition at PD was suppressed in leaves infiltrated with BIR1-HA, with callose levels remaining similar to H_2_O-treated controls (Figure 6a,b). Western blot analysis confirmed BIR1-HA protein accumulation in infiltrated tissue at 2 dpi (Figure 6c). This indicates that BIR1 inhibits poly(I:C)-induced callose deposition. To investigate whether BIR1 supports virus accumulation and spread, we co-expressed TRV-GFP and BIR1 (HA- or mCherry-tagged) under the 35S promoter in *N. benthamiana* leaves using Agrobacterium-mediated transient assays. Western blot analysis revealed that BIR1-HA had no effects on TRV accumulation as determined by TRV-derived GFP protein levels in the infiltrated area at 3 and 5 dpi (Figure 6d). However, our results demonstrated that at 2 dpi, TRV-derived GFP fluorescence extended significantly beyond the infiltrated area more rapidly in leaves expressing BIR1-HA or BIR1-mCherry compared to EV controls (Figure 6e). This suggests that BIR1 induction facilitates TRV cell-to-cell movement, but not TRV proliferation, during infection in *N. benthamiana*.

**Figure 6.**
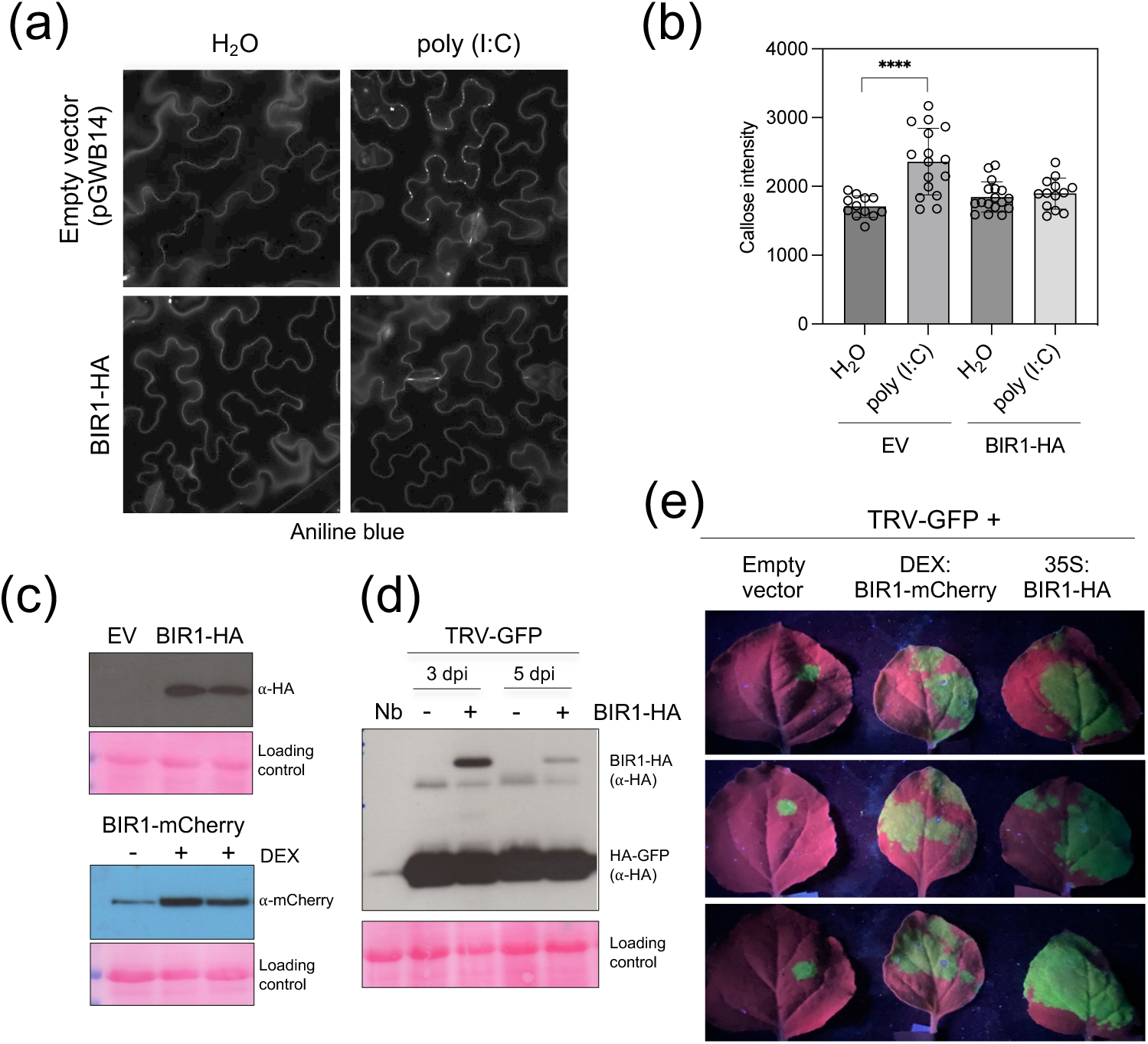
Transient BIR1 expression partially inhibits poly(I:C)-triggered callose deposition at plasmodesmata (PD) in *N. benthamiana*. (a) Callose deposition at PDs was visualized by aniline blue staining of epidermal cells in *N. benthamiana* leaves transiently expressing a 35S-driven BIR1-HA construct or an empty vector (EV). Callose deposition was observed after treatment with H_2_O (control) or 50 ng/µl poly(I:C). (b) Quantification of PD callose content in leaves infiltrated with EV or BIR1-HA in response to H_2_O or poly(I:C). Each dot represents the mean callose intensity at individual PDs (>100) measured in three to four leaf discs from at least three independent biological replicates per treatment. Statistical significance was determined by one-way ANOVA followed by Tukey’s multiple comparison test; ****adjusted p-value < 0.0001; ***p < 0.001; **p < 0.01. (c) Western blot analysis using anti-HA and anti-mCherry antibodies to detect BIR1-HA and BIR1-mCherry proteins, respectively, in agroinfiltrated areas at two days post-infiltration (dpi). (d) Western blot analysis using anti-HA antibodies to detect HA-tagged BIR1 and GFP proteins in areas co-infiltrated with TRV-GFP and EV (-) or BIR1-HA (+) at three- and five-days post-infiltration (dpi). Red Ponceau is shown as loading control. (e) Transient expression of BIR1 promotes viral spreading in *N. benthamiana* leaves. Leaves were co-infiltrated with a mixture of Agrobacterium containing TRV1 and TRV2-GFP (1:1) plus BIR1-HA under the 35S promoter, BIR1-mCherry under a dexamethasone (DEX)-inducible promoter, or the corresponding empty vector (EV). For DEX-inducible BIR1-mCherry expression, 30 µM DEX was added to the infiltration solution. Photographs were taken under UV light two days post-infiltration to visualize GFP fluorescence. All experiments were repeated three times with similar results.

### BIR1 induction hinders PD callose deposition during PTI in Arabidopsis

We next investigated whether induction of BIR1 affects PD callose deposition using BIR1 L9 Arabidopsis transgenic plants. Expression of mCherry-tagged BIR1 was induced with DEX, while control plants were mock-treated with H_2_O. In WT plants, poly(I:C) treatment significantly increased PD-associated callose levels compared to H_2_O treatment, regardless of DEX application, confirming that DEX alone does not affect dsRNA-induced callose deposition at PD (Figure 7a,b). Similarly, poly(I:C) elicited PD callose in naïve BIR1 L9 plants (without DEX) compared to H_2_O-treated controls (Figure 7c). However, BIR1 induction via DEX significantly suppressed poly(I:C)-induced PD callose deposition, which remained at H_2_O-induced levels (Figure 7c,d).

**Figure 7.**
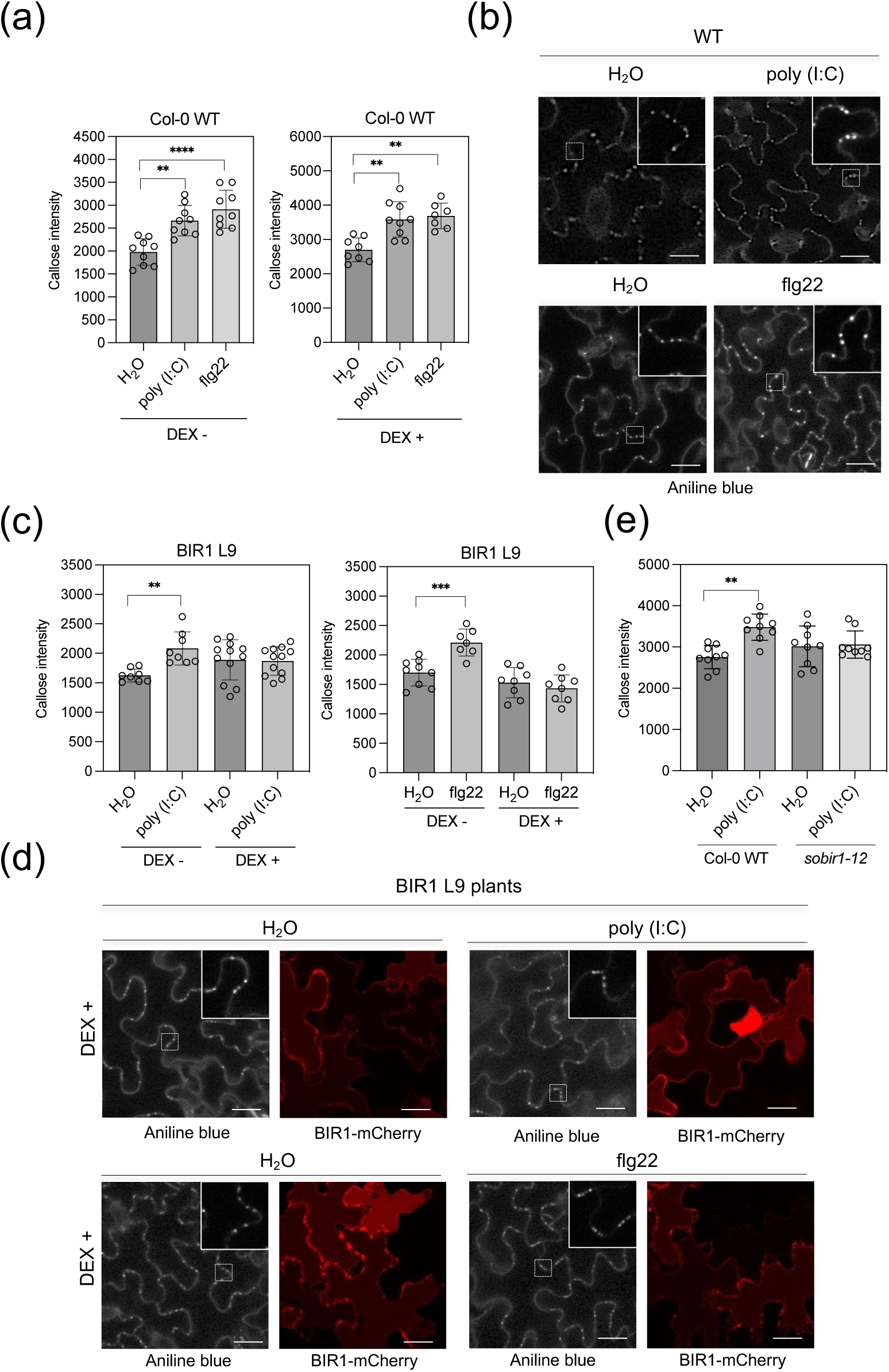
BIR1 induction inhibits flg22- and poly(I:C)-triggered callose deposition at plasmodesmata (PD) in Arabidopsis. (a) Quantification of PD callose content in wild-type (WT) plants pre-treated with or without 30 µM dexamethasone (DEX) for three days in response to H_2_O (control), 50 ng/µl poly(I:C), or 1 µM flg22. (b) Aniline blue staining to visualize callose deposition at PDs in epidermal cells of WT plants in response to poly(I:C) or flg22. (c) Quantification of PD callose content in BIR1 L9 plants pre-treated with or without DEX in response to H_2_O, poly(I:C), or flg22. (d) Aniline blue staining of BIR1 L9 transgenic plants pre-treated with or without DEX in response to H_2_O, poly(I:C), or flg22. Photographs were taken 30 minutes after treatments. BIR1-mCherry fluorescence was confirmed in DEX-treated BIR1 L9 plants within the same images. Inlays show magnified views of the boxed areas. Scale bar: 10 µm. (e) Quantification of PD callose content in WT and *sobir1-12* mutant plants in response to H_2_O or poly(I:C). (a,c,e) Each dot represents the mean callose intensity at individual PDs (>100) measured in three to four leaf discs from at least three independent biological replicates per treatment. Statistical significance was determined by one-way ANOVA followed by Tukey’s multiple comparison test; ****adjusted p-value < 0.0001; ***p < 0.001; **p < 0.01. All experiments were repeated twice with similar results.

To determine if BIR1 affects PD callose deposition triggered by non-viral elicitors, we treated plants with 1 µM bacterial flg22. In WT plants, flg22 significantly increased callose deposition at PD compared to H_2_O, irrespective of DEX treatment (Figure 7a,b). In BIR1 L9 plants, the flg22-induced PD callose deposition observed in the absence of DEX was partially suppressed when BIR1 was induced with DEX (Figure 7c,d). These results suggest that BIR1 disrupts a shared step in the defense signaling pathway regulating PD trafficking across different elicitors. Additionally, BIR1-mediated suppression of PD callose deposition appears conserved in *N. benthamiana* and Arabidopsis.

To evaluate whether PD callose deposition is a resistance mechanism to TRV in *bir1-1* mutants, we examined dsRNA-induced PD callose levels in *sobir1-12* mutants. Poly(I:C) triggered callose deposition in WT plants but not in *sobir1-12* mutants, where PD callose levels after poly(I:C) treatment were similar to those with H_2_O (Figure 7e). This indicates that PD callose deposition triggered by poly (I:C) involves the SOBIR1 co-receptor and suggests a secondary role for callose deposition in the SOBIR1-independent antiviral resistance observed in *bir1-1* mutants.

### BIR3 regulates BAK1- and EDS1-dependent antiviral resistance in Arabidopsis

We investigated the role of BIR2 and BIR3 in antiviral defense using the *bir2-1* and *bir3-2* T-DNA insertion mutants (Halter *et al*., 2014; Imkampe *et al*., 2017), which exhibit WT morphologies and are suitable for TRV inoculation. Like BIR1, RT-qPCR analysis revealed that *BIR2* and *BIR3* depletion significantly reduced TRV accumulation at 5 dpi, indicating their negative regulation of antiviral signaling pathways (Figures 2a, 8a). We then examined the role of immune components downstream of BIR3 in TRV resistance in *bir3-2* mutants. The intracellular TNL receptor *CONSTITUTIVE SHADE AVOIDANCE 1* (*CSA1*) senses integrity of BIR3-BAK1 complexes in uninfected plants, and activate EDS1/PAD4-dependent cell death pathways in *bir3-2 bak1-4* mutants (Schulze *et al*., 2022). Severe dwarfism and spontaneous cell death in the *bir3-2 bak1-4* mutant rendered TRV inoculation impossible, and excluded it from our analysis. However, the *bir3-2 bak1-4 eds1-12* mutant, where the *eds1-12* mutation partially suppresses *bir3-2 bak1-4* phenotypes, accumulated TRV at WT levels, suggesting that *BAK1* and *EDS1* mutations nullify the antiviral response in *bir3-2* plants (Figure 8a). Notably, the dwarf phenotype of *bir3-2 bak1-4* mutants is also restored by a T-DNA insertion in the *CSA1* gene (Schulze *et al*., 2022). TRV accumulation in the *csa1-2* single mutant matched WT levels, showing that CSA1 does not activate antiviral defenses (Figure 8a). Similarly, TRV levels in the *bir3-2 bak1-4 csa1-2* triple mutant were comparable to WT, indicating that BAK1, but not CSA1, mutations reversed TRV resistance in *bir3-2 bak1-4 csa1-2* plants (Figure 8a). These results demonstrate that BIR3 negatively regulates BAK1- and EDS1-dependent antiviral signaling, distinct from BIR1 mechanisms.

**Figure 8.**
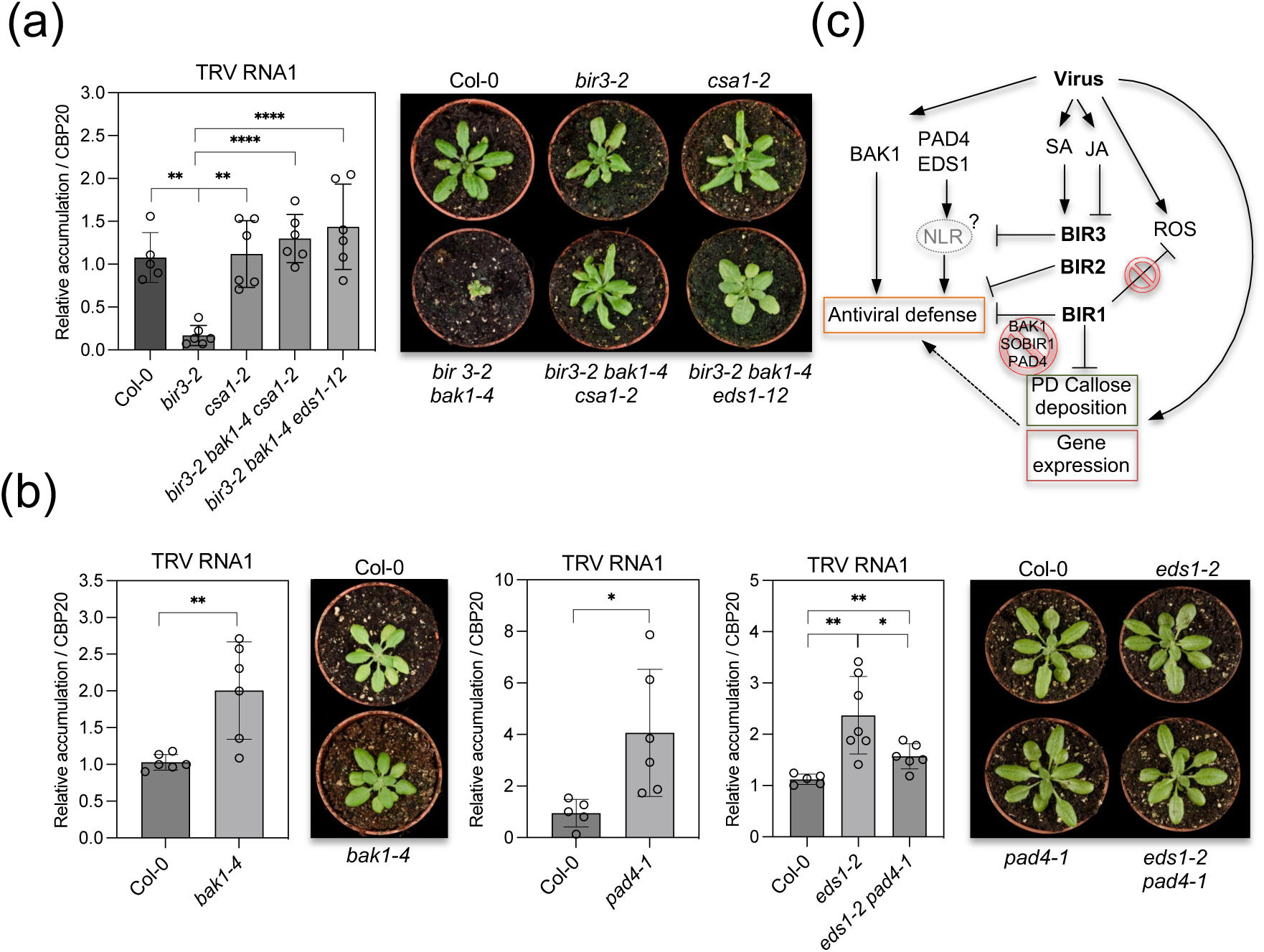
Genetic dissection of the antiviral response in *bir3-2* mutants. (a) RT-qPCR analysis of TRV genomic RNA levels in TRV-infected Arabidopsis wild-type (WT, Col-0) and various mutant combinations, including *bir3-2*, *csa1-2*, *bir3-2 bak1-4 csa1-2*, and *bir3-2 bak1-4 eds1-12*, at 5 days post-inoculation (dpi). (b) RT-qPCR analysis of TRV genomic RNA levels in TRV-infected WT (Col-0), *bak1-4, pad4-1*, *eds1-2*, and *pad4-1 eds1-2* at 5 dpi. Morphological phenotypes of each genotype at 5 dpi with TRV are also shown. A non-inoculated *bir3-2 bak1-4* mutant is included for size comparison. Relative TRV RNA levels were normalized to the *CBP20* internal control and compared to wild-type plants (set to 1). Data are presented as mean ± SD. Statistical significance was determined by one-way ANOVA followed by Tukey’s multiple comparison test; ****, p <0.0001; ***, p< 0.001; **, p< 0.01; *, p <0.05. Experiments were repeated at least twice with similar results. (c) Model for BIR-mediated regulation of antiviral defense. The BIR proteins (BIR1, BIR2, and BIR3) are induced during virus infection, influenced by antagonistic interactions between salicylic acid (SA) and jasmonic acid (JA) hormone signaling. BIR1 suppresses pattern-triggered immunity (PTI) gene expression and plasmodesmata (PD) callose deposition, contributing to antiviral defense through mechanisms that may include yet unidentified pathways independent of reactive oxygen species (ROS) or the BAK1, SOBIR1, or PAD4 signaling components. BIR3 represses the activation of the BAK1- and EDS1/PAD4-dependent effector-triggered immunity (ETI) antiviral response, likely involving intracellular nucleotide-binding leucine-rich repeat (NLR) receptors, leading to asymptomatic resistance.

To confirm the antiviral roles of BAK1 and EDS1, we inoculated *bak1-4* and *eds1-12* single mutants and measured TRV accumulation. Both mutants showed significantly higher TRV levels than WT plants at 5 dpi, corroborating their roles in antiviral resistance (Figure 8b). Previous studies also demonstrated increased TRV susceptibility in the *bak1-5* allele, further validating the contribution of BAK1 to TRV virus resistance (Guzman-Benito *et al*., 2019). We extended this analysis to PAD4, an EDS1 partner, and found elevated TRV levels in *pad4-1* single and *eds1-12 pad4-1* double mutants, highlighting the role of PAD4 in the antiviral pathway (Figure 8b). Consistent with these results, the *sobir1-1 bak1-4 pad4-1* triple mutant also exhibited heightened TRV susceptibility compared to WT (Figure 2a). Interestingly, TRV accumulation occurred across all tested genotypes without macroscopic hypersensitive response, suggesting that BAK1/EDS1/PAD4-mediated resistance operates independently of cell death (Figure 8a,b). In conclusion, BIR1, BIR2, and BIR3 negatively regulate at least two independent defense pathways that restrict viral accumulation in Arabidopsis, underscoring the conserved role of the BIR family as modulators of antiviral defense.

## Discussion

Plants utilize a two-layered immune system to combat microbes like bacteria, fungi, and oomycetes. Recent studies reveal that canonical immune responses, including PTI and ETI, also influence plant-virus interactions (Calil & Fontes, 2017; Piau & Schmitt-Keichinger, 2023). While some (co)receptors and immune pathways that defend against non-viral pathogens affect viral resistance and susceptibility, the mechanisms by which specific PTI and ETI antiviral responses are initiated and transduced remain unclear (Macho & Lozano-Duran, 2019). A significant breakthrough was the discovery that viral dsRNA activates a PTI signaling pathway via the pattern recognition co-receptor kinase SERK1 (Niehl *et al*., 2016). dsRNA-induced PD callose deposition also requires PTI components, such as BIK1, PBL1, and PD-localized proteins PDLP1, PDLP2, and PDLP3 (Huang *et al*., 2023). Previously, we found that *BIR1* expression is induced during virus infection, likely in an SA-dependent manner, as SA accumulates in TRV-infected plants and promotes *BIR1* transcription (Guzman-Benito *et al*., 2019). In this study, we observed that TRV infection also increased the expression of *BIR2* and *BIR3*, suggesting a role for BIR proteins during infection. Alongside the upregulation of *BIR1*, *BIR2*, and *BIR3*, TRV triggered the accumulation of the defensive hormones SA and JA. SA treatments enhanced *BIR1* and *BIR3* expression, while JA suppressed them. Although the role of JA in virus defense remains unclear, our findings suggest that the concurrent activation of SA and JA pathways during viral infection is critical for maintaining BIR signaling homeostasis by antagonistically regulating *BIR1* and *BIR3* expression in Arabidopsis (Figure 8c).

We demonstrate that BIR proteins act as negative regulators of TRV resistance, highlighting a conserved role for the BIR family in modulating antiviral defense in Arabidopsis. Our genetic evidence indicates that BIR1 and BIR3 function through distinct mechanisms. Previous studies showed that BIR1 knockout in uninfected Arabidopsis causes cell death via SOBIR1- and EDS1/PAD4-dependent pathways (Gao *et al*., 2009). This raises the question of whether these same pathways drive the antiviral resistance observed in *bir1-1* mutants, characterized by reduced TRV accumulation. Our findings reveal that the antiviral response in the *bir1-1 sobir1-1* double mutant is unaffected by the loss of the adapter kinase SOBIR1, while PAD4, a key regulator of TNL-mediated ETI and PTI, is not required for TRV resistance in *bir1-1* mutants. Additionally, although BAK1 contributes to *bir1-1* phenotypes (Liu *et al*., 2016), it is not necessary for *bir1-1*-mediated TRV resistance. Overall, our results indicate that the immune pathways driving cell death, constitutive defense activation, and disease resistance in *bir1-1* mutants are distinct from the antiviral response against TRV that is triggered by BIR1 disruption (Figure 8c).

We report that induced BIR1 expression negatively regulates the transcription of several flg22-responsive PTI marker genes when the PTI elicitor flg22 is present. This is significant because BIR1 induction alone does not affect defense gene expression, indicating that BIR1 specifically interferes with immune gene transcription during PTI activation. Additionally, the antiviral response in *bir1-1* mutants appears to be unrelated to ROS, similar to the dsRNA-induced PTI pathway that triggers PD callose deposition independently of ROS (Huang *et al*., 2023). This prompted us to explore the effect of BIR1 induction on dsRNA-triggered callose deposition at PD. PD callose deposition restricts virus movement in plants, with viruses countering this defense by encoding suppressor proteins such as tobamovirus MP (Huang *et al*., 2023). We found that induced BIR1 expression in both *N. benthamiana* and Arabidopsis suppresses callose deposition at PD elicited by poly(I:C) or flg22. This suggests that BIR1 induction during viral infection promotes virus spread. Supporting this, transient BIR1 expression in *N. benthamiana* facilitates TRV movement beyond the infiltration site, though further confirmation is needed in Arabidopsis. Interestingly, the antiviral response in *bir1-1* mutants is independent of BAK1, aligning with the observation that dsRNA-induced PTI-like responses depend on SERK1, not BAK1/SERK3 or BKK1/SERK4 (Niehl *et al*., 2016; Niehl & Heinlein, 2019; Huang *et al*., 2023). These findings suggest that dsRNA acts as a bona fide PTI elicitor in TRV-infected plants, with callose deposition at PD representing an antiviral mechanism downstream of BIR1. However, the antiviral response in *bir1-1* mutants does not require SOBIR1, whereas dsRNA-induced callose deposition is SOBIR1-dependent. This implies that callose deposition, while potentially contributing to *bir1-1* resistance by possibly restricting cell-to-cell virus movement, is not the sole mechanism limiting viral proliferation. The antiviral immunity in *bir1-1* mutants shares features with classic PTI responses induced by microbial elicitors but involves distinct signaling components, lacking PTI hallmarks like BAK1 signaling and apoplastic ROS production (Figure 8c) (DeFalco & Zipfel, 2021). Furthermore, our study does not conclusively establish dsRNA as a trigger for antiviral resistance linked to BIR1 function. This highlights the need for further research to unravel additional defense pathways in *bir1-1*-mediated viral resistance.

Genetic defects in BAK1 reversed TRV resistance in *bir3-2* mutants, highlighting BAK1 as a key regulator of antiviral defense in the absence of BIR3. This suggests that BIR3 may inhibit the role of BAK1 in promoting antiviral PTI, consistent with the broader function of BIR proteins as negative regulators of BAK1-dependent PTI responses (Gao *et al*., 2009; Halter *et al*., 2014; Imkampe *et al*., 2017; Wu *et al*., 2020). While this study and others confirm a positive role of BAK1 in viral defense (Macho & Lozano-Duran, 2019), it remains unclear whether this role is directly linked to BIR3-mediated regulation or involves distinct activation mechanisms—a topic for future research. The genetic basis underlying antiviral pathways activated in *bir2* mutants also remains unexplored. Notably, BAK1 has been identified as a convergence point for PTI and ETI pathways (Schulze *et al*., 2022), raising the question of whether its antiviral role depends on PTI, ETI, or both. Intriguingly, BAK1 interacts with the transmembrane receptor-like kinase NIK1 (Smakowska-Luzan *et al*., 2018), which regulates antiviral defense against Begomovirus. Upon virus infection, NIK1 represses translational machinery-associated genes, reducing viral and host mRNA translation (Carvalho *et al*., 2008; Zorzatto *et al*., 2015). This raises the possibility that BAK1 contributes to antiviral defense via NIK1 rather than through conventional PTI or ETI (Macho & Lozano-Duran, 2019).

Our reverse genetic analyses identified EDS1 and PAD4 as key positive regulators of antiviral defense. The antiviral role of EDS1 was further demonstrated in *bir3-2* mutants. Notably, the observation that TRV accumulated to similar levels in *bir3-2 bak1-4 csa1-2* and *bir3-2 bak1-4 eds1-12* mutants suggests that BAK1 and EDS1 function within the same resistance pathway. EDS1 and PAD4 are known to transduce signals downstream of NLR proteins. While the NLR CSA1 is critical for triggering cell death in *bak1-4 bir3-2* and *bak1-3 bkk1* mutants (Schulze *et al*., 2022), it appears dispensable for the antiviral defense activated by BIR3 loss. Although our data excludes CSA1, consistent with the guarding hypothesis for BIR3-BAK1 homeostasis (Schulze *et al*., 2022), we hypothesize that viruses disrupt BIR3-BAK1 complexes, triggering an antiviral EDS1/PAD4-dependent ETI response mediated by unidentified NLRs. In this model, ETI activation relies on sensing virus- and/or viral effector-induced host protein modifications rather than direct effector recognition. Notably, ETI against TRV confers symptomless resistance without triggering cell death. We propose that, in Arabidopsis, BAK1- and/or EDS1/PAD4-dependent pathways downstream of BIR3 restricts viral proliferation while allowing systemic virus movement, thereby preserving the integrity of virus-infected cells (Figure 8c).

In conclusion, our study advances the understanding of signal transduction pathways in plant-virus immunity. We show that the RLKs BIR1 and BIR3 act as facilitators of infection by dampening antiviral defenses through distinct mechanisms. BIR1 restricts PD callose deposition and PTI gene expression, although antiviral resistance in *bir1-1* mutants likely involves additional yet unknown defense pathways that limit viral proliferation. BIR3, potentially with BAK1, modulates antiviral PTI and ETI responses involving EDS1/PAD4 signaling, revealing a novel, symptomless antiviral mechanism. One question concerns the possible implications that could arise from the activation of BIR proteins during viral infections, given that they at least partially compromise antiviral defenses. In a susceptible plant, infection results from the balance between defense activation and suppression. In this scenario, BIR proteins may act as proviral factors, reducing the efficiency of PTI and ETI responses. Thus, BIR proteins may help maintain a balanced relationship that permits infection, providing beneficial effects on plant fitness without causing host damage. For example, TRV infection is asymptomatic in Arabidopsis and *N. benthamiana*, and enhances abiotic stress resistance potentially improving fitness (Fernandez-Calvino *et al*., 2014). Future research should investigate this topic further and determine whether these findings extend to other plant-virus systems

## Acknowledgements

We thank members of the César Llave lab, the Manfred Heinlein lab and the Thorsten Nürnberger and Birgit Kemmerling labs for helpful discussions. We also thank Monica Fontenla (Centro de Investigaciones Biológicas Margarita Salas, CSIC, Madrid, Spain) for her photographic work and Sonia Osorio and José Vallarino (Instituto de Hortifruticultura Subtropical y Mediterránea, UMA-CSIC, Málaga, Spain) for hormone analysis.

## Author contributions

CL conceived and designed the study. CL, CR and IGB, performed gene expression and infection assays in Arabidopsis. CL, CH, CR and MC performed ROS and ethylene assays. CL, ARS and LEG performed callose and infection assays. CL, CR, MH and TN analyzed the data. CL wrote the article.

## Supplemental data

**Figure S1.** Transcriptomic profiling of dexamethasone (DEX)-treated wild-type (WT) Col-0 and BIR1 line 9 (L9) plants. (a) Scatterplot of selected gene ontology (GO) terms and (b) Kyoto Encyclopedia of Genes and Genomes (KEGG) pathways using RNA-Seq data. Dot size represents the count of different genes and the color indicates significance of the term enrichment. Terms with adjusted *p*-value < 0.05 are significantly enriched. WT, wild-type Col-0; De, DEX-treated plants; Mo, Mock-treated plants

**Table S1.** List of primers

**Table S2.** List of expressed genes.

**Table S3.** Pearson correlation coefficient between samples for each condition.

**Table S4.** List of differentially expressed genes (DEG. |Log_2_fold change > 1| padjust value < 0.05).

**Table S5.** List of differentially expressed genes (DEG. |Log_2_fold change > 1| padjust value < 0.01).

**Table S6.** Enrichment analysis of Gene Ontology (GO) terms.

**Table S7.** Kyoto Encyclopedia of Genes and Genomes (KEGG) pathway enrichment analysis.

## Data availability

The data supporting this study’s findings are available from the corresponding author upon reasonable request

